# Effects of Connectivity on Narrative Temporal Processing in Structured Reservoir Computing

**DOI:** 10.1101/2022.08.11.503616

**Authors:** Peter Ford Dominey, Timothy M. Ellmore, Jocelyne Ventre-Dominey

## Abstract

Computational models of language are having an increasing impact in understanding the neural bases of language processing in humans. A recent model of cortical dynamics based on reservoir computing was able to account for temporal aspects of human narrative processing as revealed by fMRI. In this context the current research introduces a form of structured reservoir computing, where network dynamics are further constrained by the connectivity architecture in order to begin to explain large scale hierarchical network properties of human cortical activity during narrative comprehension. Cortical processing takes place at different time scales depending on the position in a “hierarchy” from posterior sensory input areas to higher level associative frontal cortical areas. This phenomena is likely related to the cortical connectivity architecture. Recent studies have identified heterogeneity in this posterior-anterior hierarchy, with certain frontal associative areas displaying a faster narrative integration response than much more posterior areas. We hypothesize that these discontinuities can be due to white matter connectivity that would create shortcuts from fast sensory areas to distant frontal areas. To test this hypothesis, we analysed the white matter connectivity of these areas and discovered clear connectivity patterns in accord with our hypotheses. Based on these observations we performed simulations using reservoir networks with connectivity patterns structured with an exponential distance rule, yielding the sensory-associative hierarchy. We then introduce connectivity short-cuts corresponding to those observed in human anatomy, resulting in frontal areas with unusually fast narrative processing. Using structured reservoir computing we confirmed the hypothesis that topographic position in a cortical hierarchy can be dominated by long distance connections that can bring frontal areas closer to the sensory periphery.

## I. Introduction

We are witnessing an interesting conjuncture in developments in neural network models of language in machine learning, the computational neuroscience of language, and structured reservoir computing. Neural representations in certain classes of language models are measurably similar to neural representations in the human brain [1]. In this context, the temporal dynamics of reservoirs have been demonstrated to display human-like coding properties for narrative event structure [2]. Finally, structured reservoir computing is beginning to investigate how constraining reservoir topology based on human connectivity can produce interesting computational properties [3].

The current research addresses a dimension of language processing that has gained increasing visibility – the temporal processing of language in a cortical hierarchy [4–8]. Intuitively, sensory-driven cortical responses should be rapid and reflect the temporal structure of the input, while cortical activity underlying higher level cognitive functions that may involve reasoning and memory will have longer time-scales of processing [9]. Computational models of cortical processing that emphasize local connectivity naturally demonstrate such hierarchies, with fast processing in nodes that receive input from the periphery, and a progressive slowing in areas farther from the periphery, as activity propagates along the hierarchy [10].

However, while the majority of cortical connections are local [11], the primate (and particularly the human) brain is riddled with white matter bundles that create direct connection short-cuts between topographically distant cortical areas [12]. Here we address the effects of such topological constraints on narrative processing in the context of a new variant of reservoir computing as illustrated in Fig. 1.

**Fig. 1.**
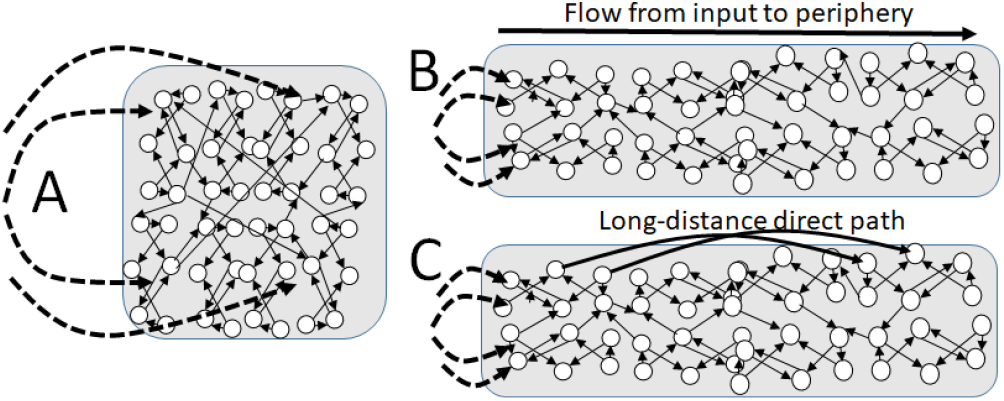
Structured Reservoir Computing. A. Standard reservoir topology, where inputs (dotted lines) are uniformly distributed into the network, and there is no restriction on the length input connections into the reservoir nor of recurrent connections (small solid lines) within the reservoir. B. Structured reservoir, where input connection are restricted to a small set of reservoir neurons, and recurrent connections are restricted by distance (i.e they are local). This creates a flow of information from input driven neurons to more distant peripheral “associative” neurons across local connections. C. Structured reservoir with direct connection short-cuts between topographically distant cortical areas, corresponding to white matter bundles.

The current research examines the relation between the different structures of network connectivity, as illustrated in Figure 1, and the resulting temporal selectivity of different cortical areas. Based on an analysis of human neurophysiology of narrative processing, different temporal processing for different cortical areas, and the underlying white matter anatomical connectivity, we develop a form of structured reservoir computing that provides a coherent explanation for these data.

### A. Temporal hierarchy in cortical processing

A temporal processing hierarchy has been observed in the human processing of narrative [4, 5]. Chien and Honey [8] demonstrated how the time scale of narrative integration increases along such a hierarchy from primary auditory cortex to the highly integrative temporal parietal junction (TPJ). They developed a task and analysis that we refer to as the Narrative Alignment Task. While recording fMRI they exposed separate subject groups to an intact narrative and a scrambled narrative. From time to time the two groups thus heard the same narrative segment. The authors compared brain activity across the two groups as they made the transition *from* listening to different narrative segments *to* listening to the same narrative segment. They aligned their analysis on this transition from different to same, and characterized the time constant for the alignment or correlation of the brain activity across the two groups as they began to hear the same narrative. Interestingly, they observed a hierarchy of alignment time constants, with fast time constants in areas near the sensory periphery (primary auditory cortex), medium time constants in intermediate areas, and slower time constants in high level integration areas (TPJ). This is consistent with related studies that identify temporal processing hierarchies for narrative processing.

One hypothesis to explain such a hierarchy of processing is that areas that are close to the sensory periphery will quickly be influenced directly by inputs, and thus in the Narrative Alignment Task will become aligned rapidly. In contrast, cortical areas that are far from the sensory periphery will take more time to respond to inputs which must traverse multiple cortico-cortical connections before arriving.

### B. Modeling the temporal hierarchy

Chaudri [10] examined a family of recurrent network models in order to characterize the relation between network architecture and temporal processing. He investigated this by using an exponential distance rule (EDR) to constrain connectivity, where the probability of a node being connected to its neighbor falls off exponentially, as schematized in Fig. 1B. This favors local connectivity. Information thus propagates through local connections from the input driven periphery to more distal nodes, with a progressive increase in the time constant for response along the resulting hierarchy. Interestingly, Chaudri further examined the effects of the introduction of long distance connections, as schematized in Fig. 1C, and demonstrated that connections from fast nodes near the base of the hierarchy to slower nodes near the top of the hierarchy can render the latter faster, thus producing a discontinuity in the hierarchy.

Such long distance connections can thus produce discontinuities in the hierarchy, such that an area that is more frontal may have response properties that are faster than an area that is more posterior. Indeed Chaudri and colleagues indicate that this appears to be the case in models constrained by the neuroanatomical connectivity of the non-human primate [13]. An open question concerns how these connectivity effects can influence the time-course of narrative processing.

### C. Modeling narrative context construction

A recently developed model of human narrative processing based on reservoir computing was used to simulate aspects of this narrative processing [2], modeling the Narrative Alignment Task of [8]. The model simulated narrative processing by using 100 dimensional word embeddings from Wikipedia2Vec [14] as inputs to a 1000 unit reservoir model of cortex. The activation of the reservoir, driven by narrative input in the form of word embeddings, thus served as a proxy for human brain activity during narrative listening. The model was demonstrated to display human-like representations of narrative event structure [2]. The model was also shown to display human-like neural representations in the Narrative Alignment Task.

Paired models were presented with narrative input (682 words) with intact (ABCD) and scrambled (ACBD) structure, respectively. The transitions CD and BD in inputs to the two models constitute the construction or integration context in the Narrative Alignment Task of [8]. That is, two models are initially exposed to different inputs (C) and (B), then to the same input (D). At the transition when both begin receiving the same input, one can measure the time constant for them to converge to the same activity. Thus, measuring the time constant for the two networks to converge to the same activity pattern when they both begin to receive common input (D) after having different inputs (C and B respectively) allows the authors to measure the alignment/convergence time. In this way the reservoir was demonstrated to simulate the Narrative Alignment Task.

Sorting the network units by these alignment times revealed a smooth distribution of alignment times from 4 to 50 time-steps, analogous to the distribution of alignment times observed in different cortical areas in humans in [8]. Interestingly, a linear integrator model of cortical function did not yield this distribution of alignment time constants – all units have the same time constants, so the distribution of time constants is a property of the flow of information in the recurrent connectivity within the reservoir.

Here we will use this narrative context construction time constant for examining the temporal processing of narrative structure, comparing across human subjects from [8], and in structured reservoir simulations, based on [2].

### D. Model performance in narrative comprehension

In the current research, we will analyze the temporal processing properties of the structured reservoir that is exposed to narrative. We thus focus on the activity of the reservoir neurons themselves, rather than on trained readouts. However, it is important to note that we have previously characterized the ability of trained reservoirs to perform integrative comprehension of narrative that includes making inferences about human event structure [15].

In [15] the readout was trained to produce an integrated representation of the input narrative as a vector average of the accumulated word embeddings constituting the narrative. These narrative embeddings were then compared to embeddings for words that were, or were not, related by inference to the narrative, following human comprehension experiments [16]. The model reproduced human performance by generating narrative mbeddings that were closer to the embeddings for words that were related to the narrative by inference. In addition, these trained narrative representations evolve in real-time during the word-by-word presentation of the narrative, again simulating human behavior in real-time integration of discourse [17]. Given this confirmation that the reservoir model can generate human-like performance in narrative/discourse comprehension, we now investigate the underlying representations within the reservoir. In particular we investigate how structural constraints on the reservoir topology can produce human-like temporal brain dynamics.

### E. Current research hypotheses and objectives

While the notion of a temporal processing hierarchy for narrative holds at a global level, details in certain studies indicate that along the posterior to anterior gradient there are inconsistencies, where more anterior areas have faster processing times than would be expected by their topographic position (e.g. [8]). We hypothesize that these discontinuities in the temporal hierarchy can be explained by long distance connections via white matter pathways between faster posterior areas and otherwise slower frontal areas that thus become faster. We test this hypothesis in a twofold approach:

First, we identify a set of cortical areas from [8] that reveal a discontinuity in the temporal processing hierarchy vs topographic hierarchy. We then analyze white matter tractography of these cortical areas, and demonstrate that indeed, the temporal processing discontinuity is correlated with the pattern of long distance white matter connections for these areas.

In order to demonstrate a causal role for long distance connections in modification of the temporal processing hierarchy, we perform structured reservoir computing simulations where the connectivity structure is organized according to an exponential distance rule, as schematized in Fig. 1B. We submit these networks to the Narrative Alignment Task of [8]. This reveals a clear gradient of temporal processing time constants within the reservoir, along the sensory-associative axis. We then introduce long distance connections corresponding to white matter bundles in the human cortex, as schematized in Fig. 1C, and demonstrate that these connections can produce discontinuities in the temporal processing hierarchy.

## II. Neural dynamics and network anatomy

In [8] the narrative context construction times obtained in the Narrative Alignment Task were determined for ~70 distinct cortical areas. The authors generally observed a hierarchical gradient, as sensory cortices aligned most quickly, followed by mid-level regions, while some higher-order cortical regions took more than 10 seconds to align.

### A. Discontinutities in the temporal processing heirarchy

Interestingly, there are discontinuities in the temporal processing hierarchy. Figure 2 illustrates three frontal cortical areas whose alignment times are in opposition to their position along the posterior-anterior topography, in data from [8]. That is, the most anterior of the three medial frontal areas indicated in the red box in Figure 2 is the fastest – in contradiction to the expected observation that the more frontal area is farther from the periphery, should be more integrative and thus have a longer time constant.

**Fig. 2.**
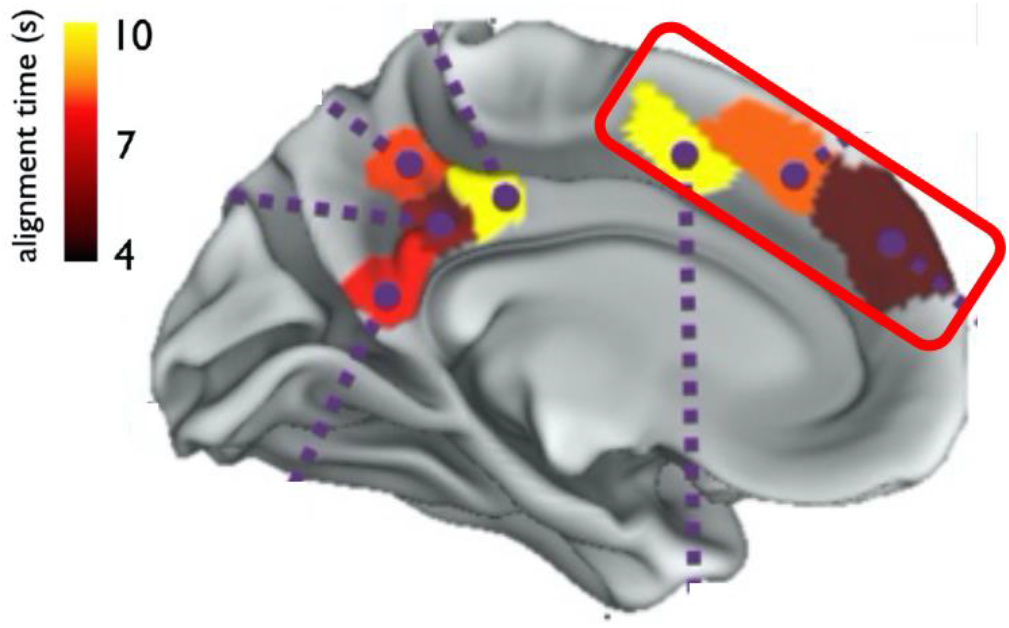
Alignment times in contrast to topography. Within the red boxed frontal zone, regions PFCd_1 (brown), PFCmp_1(orange) and FrMed_2 (yellow), lie along a topographic gradient from anterior to posterior. Their alignment times are in opposition to this topographic gradient, with the more anterior regions displaying faster alignment times. Modified from [8].

### B. Discontinuities in connectivity due to white matter bundles

In order to determine whether the temporal processing discontinuities observed by [8] for these three regions could be related to differences in white matter connectivity, we used diffusion tensor imaging [18] to characterize white matter pathways issued from these regions of interest (ROIs). Diffusion tensor data obtained from 19 subjects in [19] were analysed in order to characterize the white matter pathways issued from the three ROIs using DSI Studio (https://dsi-studio.labsolver.org).

Figure 3 illustrates white matter pathways that project from these three regions of interest (ROIs). These were generated using ROIs specified in [8], based on the atlas in [20]. The observation of a distinction between the long anterior-posterior projections for areas PFCd and PFCmp that is not seen in FrMed was confirmed by a statistical analysis of the mean lengths of the tracts issued from these three areas. Analysis of variance by repeated measure ANOVA revealed a significant effect for Region (F(2,36) = 42, p < 0.0001). Post-hoc (Scheffe) tests revealed that the mean lengths for FrMed were significantly shorter (55mm) than for PFCd and PFCmp (70, 75mm respectively), while there was no difference between the lengths for PFCd and PFCmp.

**Fig. 3.**
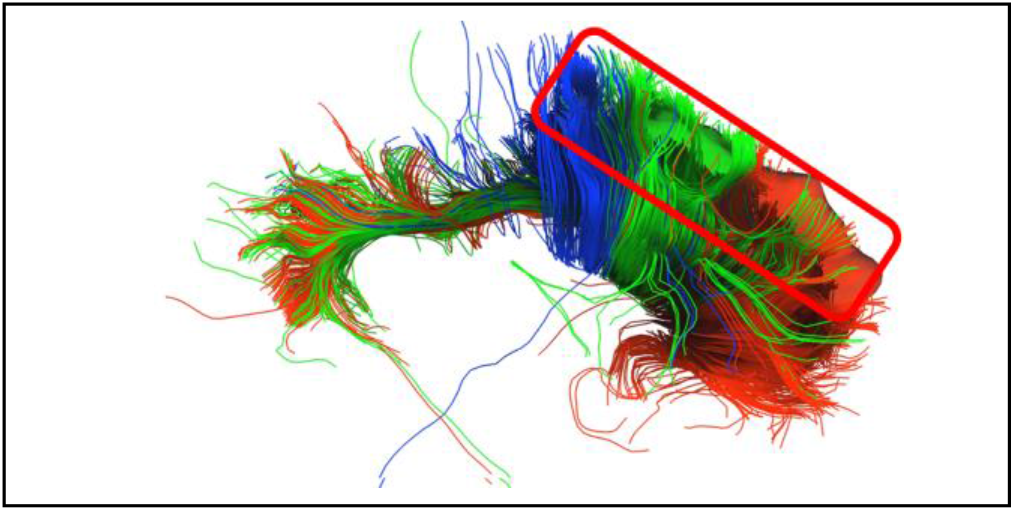
White matter connectivity for PFCd_1 (Red), PFCmp_1 (Green) and FrMed_2 (Blue). Right is anterior. Red-boxed zone same as in Fig. 2. Note that Red and Green pathways extend in a signifant manner into posterior regions, while the blue pathways have less extension into posterior regions.

Thus, the fastest area has significantly longer white matter pathways than the slowest area. Examining these pathways in Figure 2, we can observe that the pathways from PFCd and PFCmp project into posterior cortex, while those from FrMed (blue fibers) do not take this path into more posterior cortical areas.

## III. STRUCTURED RESERVOIR MODELING

These results argue in favor of the hypothesis that differences in white matter connectivity can influence the temporal processing characteristics of cortex. To test this in an empirical model of cortical dynamics we developed a structurally modified version of the narrative integration reservoir using the open access reservoir model from [2].

### A. Narrative Integration Reservoir

The classic reservoir architecture is illustrated in Fig. 1A. Our model is based on a classic echo state network with leaky integrator tanh units. A set of recurrently connected nodes – the reservoir – is stimulated by inputs. This produces a dynamic reverberation of activation throughout the reservoir as information propagates through the recurrent connections.

The basic discrete-time, tanh-unit echo state network with *N* reservoir units and *K* inputs is characterized by the state update equation:

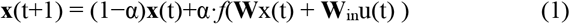

where **x**(*n*) is the *N*-dimensional reservoir state, *f* is the tanh function, **W** is the *N*×*N* reservoir weight matrix, **W***in* is the *N*×*K* input weight matrix, **u**(*n*) is the *K* dimensional input signal, α is the leaking rate. The matrix elements of W and W_in_ are drawn from a random distribution.

The reservoir was instantiated using easyesn, a python library for recurrent neural networks using echo state networks (https://pypi.org/project/easyesn/) [21]. We used a reservoir of 1000 neurons, with input and output dimensions of 100. The W and W_in matrices are initialized with uniform distribution of values from −0.5 to 0.5, with 20% non-zero connections. The leak rate was 0.2. The reservoir is relatively robust to changes in these values, as long as the reservoir dynamics are neither diverging nor collapsing.

To simulate narrative processing, words were presented in their sequential narrative order to the reservoir. Stop words (e.g. the, a, it) were removed, as they provide no semantic information [22]. Similar results were obtained in the presence of stop words. Words were coded as 100 dimensional vectors from the Wikipedia2vec language model.

This language model is developed based on using the word2vec language model that is trained on the entire Wikipedia 2018 corpus

The reservoir model illustrated in Fig. 4 receives as input the 100 dimensional word embedding from Wikipedia2Vec for the next word in the 682 word input narrative, based on the story “It’s Not the Fall that Gets You” by Andy Christie (https://themoth.org/stories/its-not-the-fall-that-gets-you). Using two identical copies of this model we can simulate the Narrative Alignment Task. One model gets the intact input narrative with segments ABCD. The second gets the scrambled narrative with segments ACBD. As illustrated in Fig 5, when the inputs are different, the activation differences are clearly visible. When the inputs become the same, then the activations progressively align and the difference goes to zero. The time constant of this alignment is the behavior of interest. Thus, it is important to note that we are analyzing the intrinsic dynamics of the reservoir itself, and not the behavior of a trained readout layer.

**Fig. 4.**
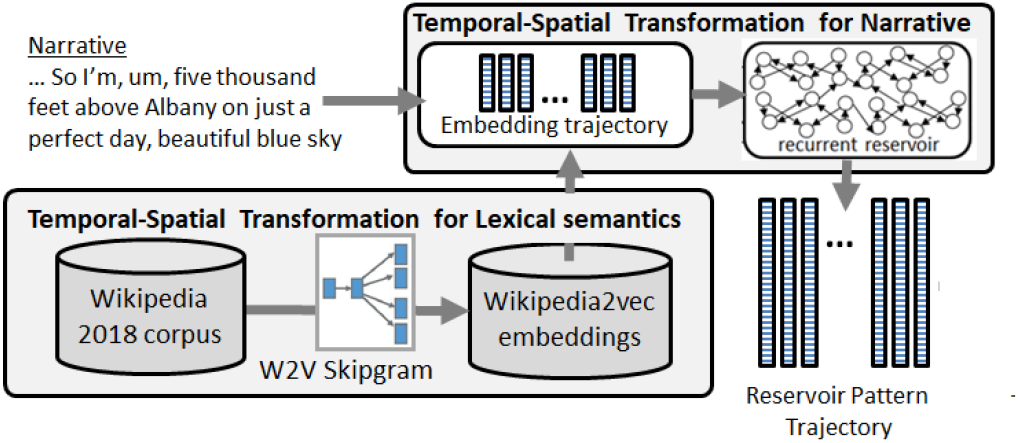
Narrative Integration Reservoir. Successive words in the input narrative are used to retreive the corresponding word embeddings from Wikipedia2vec, a word2vec model trained on the 2018 Wikipedia corpus. Successive word vectores are input to the 1000 unit reservoir model which performs a temporal-spatial transformation of temporal seqeunce of word inputs into a trajectory of spatial activation vectors.

**Fig. 5.**
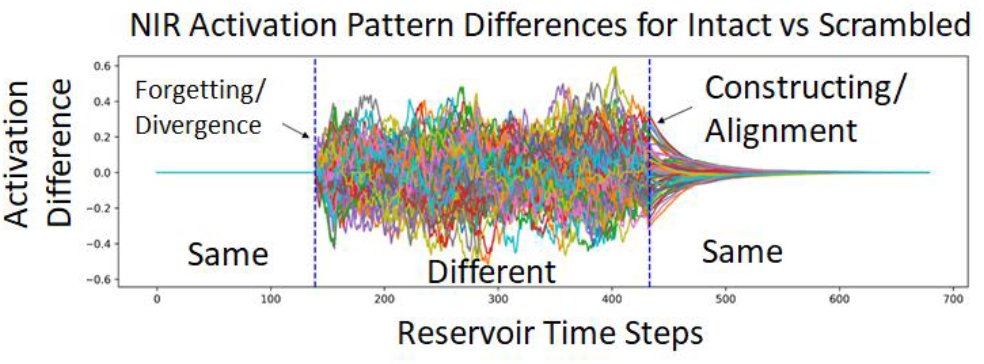
Simulation of Narrative Alignment Task. Difference in reservoir neural activity for two reservoirs exposed to intact and scrambled narratives respectively. At the transition from Different to Same, we measure the time constant for the two reservoirs to become aligned for each of 1000 neurons.

Figure 5 illustrates the difference between reservoir activity in two identical copies of the same reservoir that are exposed to the intact ABCD narrative vs. the shifted ACBD narrative. When the activations in the two reservoirs are identical, the difference is null. In the beginning the intact and shifted narratives are the same, which yields a zero difference. Then they become different and we observe the large activity in the difference signal. Finally, the two narratives again become the same. There, we see a gradual reduction in the difference as the two narratives converge to the same activity, based on the same input. The time constant of this convergence is the measure we use to characterize temporal processing of narrative based on the same measure in human subjects in [8].

### B. Topography of Temporal Processing

In the classic reservoir, the connections in the recurrent network – the reservoir – are randomly initialized with a Gaussian distribution between −0.5 and 0.5 with 20% non-zero connections. Thus, there is no structured topography within the reservoir. This is schematically illustrated in Figure 1A. In particular, there is no influence of topographical distance between neurons influencing the probability of connections.

Such a reservoir configuration, and its behavior in the Narrative Alignment Task is illustrated in Figure 6.

**Fig. 6.**
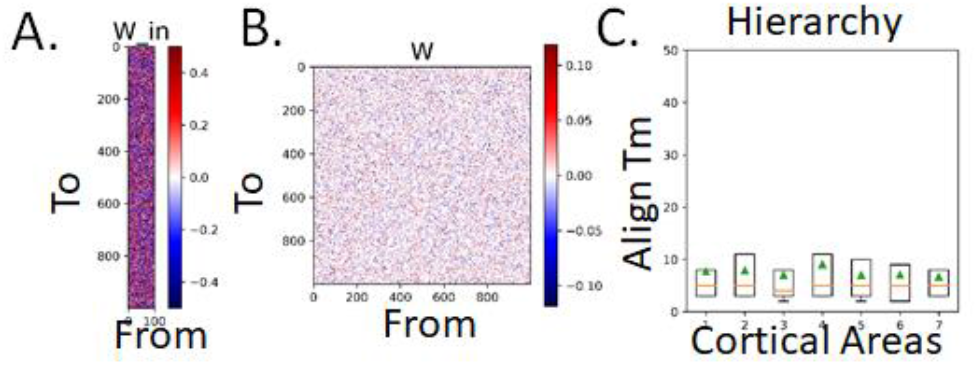
Classic reservoir architecture and “flat” temporal processing hierarchy in the Narrative Alignment Task. A. Input matrix: 100 input elements project into 1000 reservoir units. B. Recurrent connectivity matrix of the reservoir. Gaussian distribution, no structural topography. C. Narritive Alignment time constants for neuron subsets 300-399, 400-499,… 900-999. Note uniformity of time constants across these “cortical areas”.

Figure 6 illustrates a standard, random reservoir, where the input matrix projects the inputs over the whole reservoir, and where there is no length restriction on connections between reservoir units, corresponding to Fig. A. There, in Panel C we see that across the different subsets of neurons there is a uniformity of time constants across the different subsets of cortical neurons. In the primate brain, this is not the case. The probability of two neurons being connected fall off exponentially with the distance between them [23]. This is illustrated schematically in Figure 1B. In structured reservoir computing (S-RC) we can introduce structure constraints such as the exponential distance rule, and restriction of input to “sensory” areas, corresponding to the schematic in Figure 1B.

To implement this, we start with a standard reservoir connection matrix W, and then restricted it according to a exponential distance rule as described in the pseudo-code in Fig. 7. There, for neurons 1..1000, no connection can be longer than 600, and those connections are scaled by an exponential factor of the length. In addition the connections are scaled by a proximal to distal factor, that increases with the distance of the source (From) neuron from the input.

**Fig. 7.**
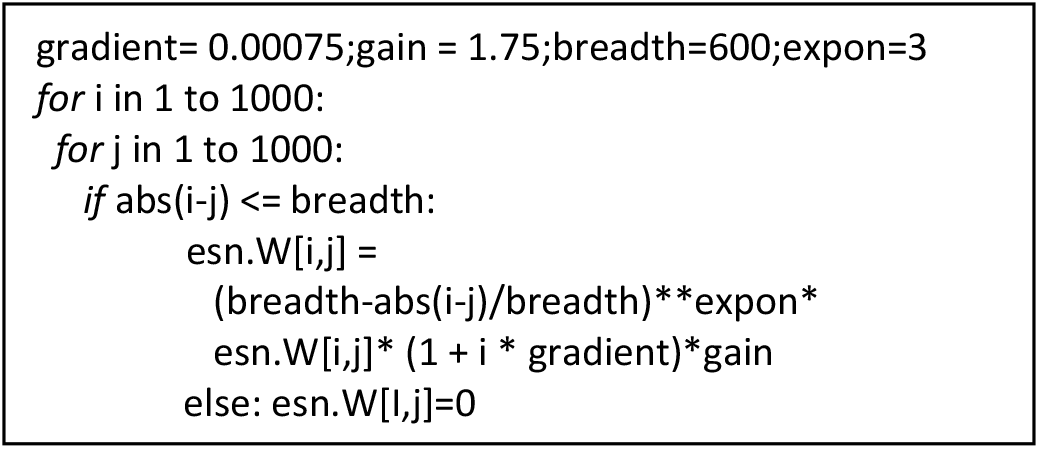
Pseudocode for creating the structured weight matrix for the recurrent reservoir connections.

In Fig. 8 we see the resulting restricted input matrix, and the structured connectivity matrix for the reservoir with an exponential distance rule applied. Values closest to the diagonal represent connections strengths between closely neighboring neurons. Values that are farther from the diagonal are for neurons that are farther apart. We thus see a density along the diagonal that falls off exponentially.

**Fig. 8.**
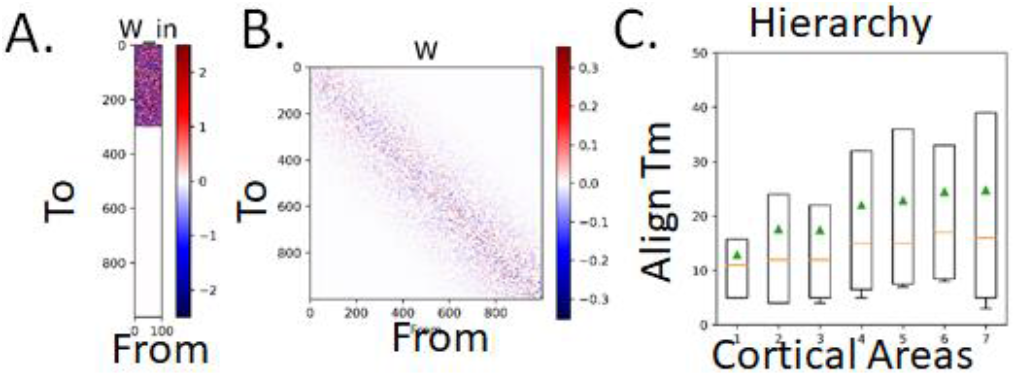
Canonical topography of temporal processing. A. In the network of 1000 neurons, inputs are provided to the the first 300 neurons. B. The connectivity matrix indicates that neurons are connected to their local neighbors according to the EDR. C. Alignment time constant based on the Narrative Alignment Task in [8].

As illustrated in Fig. 8, the EDR connectivity imposes a topographical structure on processing. While inputs directly influence “lower” areas (neurons 1-300), higher areas receive input-related activity only as it is propagated along local connections. The highest neurons will receive input-related activity only after it has traversed multiple intermediate local connections to finally arrive at the highest neurons. Panel C illustrates the functional consequences of this. Neurons are grouped into virtual areas of 100 neurons, and the mean Narrative Alignment Task time constants are displayed. We observe a progressive increase in the alignment time constants along the cortical hierarchy.

### C. Impact of Long Distance Dense Connections

This corresponds to a brain model that strictly follows local connectivity imposed by an exponential distance rule. However, recalling Figure 2 we know that thick bundles of axons provide dense connections between distant cortical areas. This corresponds to white matter – myelinated neural axons – the wire of the brain – which makes up almost half of the volume of the human brain [12]. Thus in addition to local connectivity there is ample possibility for long distance “hotlines” between cortical areas. In order to simulate the effects of such a long distance connection we introduced a connection bundle from fast input driven neurons to slower neurons far away in the hierarchy. This is illustrated in Fig. 8 panel B showing the W connectivity matrix, as the small square of values at From 100-200 to To 800-900. That is, this simulated white matter pathway connects neurons 100-200 which receive direct sensory input, to neurons 800-900 which are far from the sensory periphery. These units normally have a relatively high time constant in the absence of the long distance connection as illustrated in Fig. 8C.

As illustrated in Figure 9, the introduction of this connection produces a significant speedup in the time constant for the Narrative Alignment Task for the cortical area that receives input from the area lower in the hierarchy. This demonstrates that introduction of long distance connections can significantly modify the time constant for a given cortical area in the Narrative Alignment Task.

**Fig. 9.**
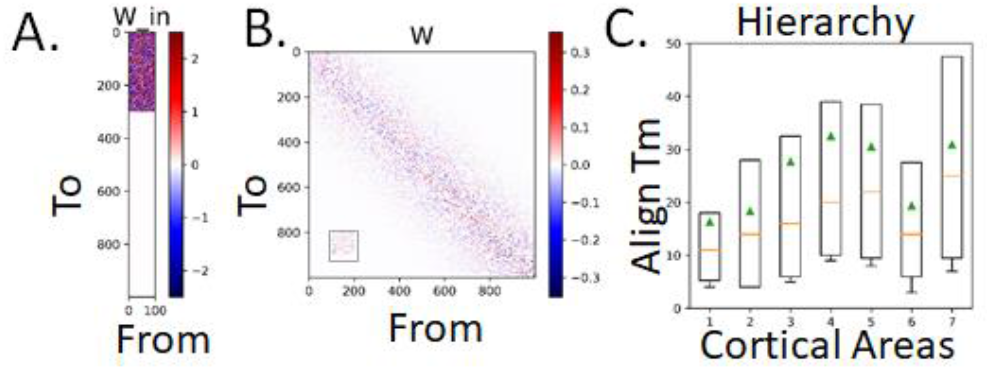
Modification of the temporal heirarchy by introduction of long distance hot-line connections. A. Input matrix. Unchanged from Fig. 7. B. Connectivity matrix W. In addition to the local EDR defined connections (seen along the diagonal) we introduce an additional white matter pathway as a direct connection from neurons 100-200 to neurons 800-900. C. Modified hierachy. Comparing to Fig 7 C, we see that that later area 6 now has a significanly reduced alignment time in the narrative alignment task.

Code and data for running these simulations is available in the repository:

github.com/pfdominey/StructuredReservoirComputing

## IV. INTERPRETATION AND DISCUSSION

The current research has scientific impact along two related dimensions. The first is the further development of a new field of investigation of structured reservoir computing where the structure of the input and recurrent connections are systematically modified based on anatomical constraints in order to produce new computational properties [3]. The second is within the context of computational neuroscience of language and narrative processing [2], and particularly the temporal component of this processing. Interestingly, these two dimensions intersect, as the analysis of computational properties of human-like connectivity in structured reservoirs will shed new light on the computational neuroscience of human narrative processing.

### A. Structured reservoir computing

Recent research in structured reservoir computing addressed how constraining the reservoir with small world connectivity enhances certain computational properties [3]. There, long distance connections allowed a propagation from restricted input-receiving units to distant output units. In their analysis, while the authors addressed the effects of the small world reservoir on several important aspects of reservoir computing, including the propagation effects of long distance shortcuts, effects on temporal processing and information flow were not directly addressed.

A clear physical phenomena is that network topology influences temporal processing [10]. When information flows from input driven areas to more distant areas, as schematized in Fig 1B, that flow takes time. We can refer to this flow of information across successive connections as the network distance. This network distance is roughly correlated with topographic or physical distance along a flattened cortical sheet. Such a network would give the flowing temporal processing properties as illustrated in Fig 8. However, the human brain is riddled with white matter (almost half the brain volume) [12]. White matter is constituted of axonal bundles which form cables that traverse the network and completely dissociate network distance from topographic distance. This is schematically illustrated in Fig. 1C. Such shortcuts should influence the temporal processing properties of their target cortical areas. In the current research this new perspective on reservoir computing is highly relevant, not only as a new avenue of reservoir computation, but also in providing a method to understand complex aspects of the human neuroscience of narrative processing.

Indeed we were motivated by a clear case in human neurophysiology where network distance and topographical distance do not correspond, based on data in [8]. That is, a more frontal area had a much faster time constant for narrative processing than more posterior neighboring areas. We hypothesized that this discontinuity could be due to different connectivity profiles for these areas. We performed DTI analysis to determine that indeed, the most frontal area which most deviated from the temporal processing according to topographical distance had a connectivity pattern that significantly varied from its neighbor that had a better correspondence between topographic and network distance (i.e. it is far from the sensory periphery, and relatively slow). In particular the faster area had more extensive connections into posterior cortical areas that are closer to the sensory periphery.

To further test the hypothesis that such long distance connections can produce speedup in temporal processing, we performed neural network simulations using a novel form of reservoir computing that we refer to as Structured Reservoir Computing.

In structured reservoir computing, we introduce structure in the matrices that describe connections from the input to the reservoir, and the recurrent connections, following related research that has investigated the effects of small-world connectivity on reservoir processing [3]. In particular, we create a more physiologically realistic reservoir topology, basing connectivity on an exponential distance rule, while retaining the reservoir principle of non-modifiable recurrent connections that are used to generate high-dimensional spatio-temporal projections of the inputs. By employing a connectivity structure where connection strength is based on an exponential function of the connection distance, we can produce structured reservoirs that display a more physiologically realistic spatial distribution of processing time constants such as observed in humans. Likewise, by introducing additional structure in the form of long distance white matter pathways that in a sense violate the exponential distance rules, we can achieve increasing realistic modeling of temporal processing as observed in [8].

An interesting and prevalent phenomena in the human processing of narrative is that across different brain areas, there are differences in the temporal granularity of narrative integration and event processing [4–8]. It will be of great interest to use structured reservoir computing to model this diversity of cortical processing, including the processing of meaning at different structural levels (e.g. word, sentence, paragraph, narrative pattern, etc.). An initial step in this direction was taken in [2]. There we demonstrated that within a classic reservoir, there was a broad distribution of narrative processing time constants as revealed in the Narrative Alignment Task. When the reservoir neurons were sorted by their alignment time constants, the distribution of alignment time constants was strikingly similar to that observed across wide distribution of cortical areas in humans [8]. In the current research we demonstrate how such a distribution of alignment time constants can be ecologically produced by imposing the exponential distance rule structure on the reservoir connectivity topology.

### B. Temporal hierarchy in narrative processing

While cortical neurons have similar intrinsic time constants, due to their similar geometry and membrane properties, there is a vast diversity of time constants of neural processing across the cortical sheet [13]. This is particularly present in narrative processing [4–8]. A major open question in modern neuroscience concerns both the origin of this diversity, and its functional consequences. One hypothesis would hold that specific cognitive processes, such as integration of narrative structure over an extended portion of narrative input, requires a long time constant of processing, and so these high level functions will impose this temporal structure on their associated brain areas or networks that are involved in such processing. In this hypothesis, the question remains as to what is the origin of the function that imposes these temporal constraints. An alternative hypothesis holds that the gross temporal processing constraints are imposed by architecture itself. As we observe, areas far from the sensory periphery naturally have slower time constants. In this hypothesis, these areas will naturally be adapted to processes that involve long time scale integration. Thus, long time scale structure of narrative will naturally self-allocate to these regions.

This modeling work examines the emergence of a temporal processing hierarchy based on network topology. It provides evidence that the distribution of narrative construction time constants observed in [8] might be accounted for by cortical network structure, rather than by an adaptation of different cortical areas to different processing requirements. In other words, the different narrative integration processes (that operate e.g. at the word, sentence, paragraph, chapter levels) naturally migrate to the brain regions that have the corresponding connectivity structure that yields the appropriate processing time constant. This is in opposition to the notion that different brain areas adapt their processing time constants for different tasks. This is a rather binary analysis. It is possible that the truth is a compromise, whereby processing time constants can be constrained both by network connectivity properties, and by on-line modulatory processes. This provides a rich framework for future research on the interaction between network structure, processing time constant dynamics modulated by input driven and intrinsic processes in biological and artificial networks [24].

On a final note, this research illustrates the tight potential link between the topological connectivity structure of the brain, and the hierarchical structure of narrative. This provides a novel and potentially fruitful approach for investigating the co-evolution of human narrative, including its spatio-temporal complexity and hierarchical event structure, and the corresponding neural architecture that provides the substrate for creating and comprehending this narrative structure. In the context of reservoir computing this is quite exiting. We already have the example of how reservoir computing has helped to characterize and understand the neuroscientific importance of mixed selectivity for higher cognitive function [25, 26]. This new research in structured reservoir computing and narrative opens a new avenue of research that one can hope will be equally productive.

## ACKNOWLEDGMENT

Reservoir simulations were performed using the easyesn toolkit https://github.com/kalekiu/easyesn.

